# Plasma irradiation efficiently inactivates the coronaviruses mouse hepatitis virus and SARS-CoV-2

**DOI:** 10.1101/2020.11.13.381319

**Authors:** Shigeru Morikawa, Shunpei Watanabe, Hikaru Fujii, Toshio Tanaka, Junichirou Arai, Shigeru Kyuwa

## Abstract

Many inactivation methods have been shown to inactivate SARS-CoV-2 for safe and efficient diagnostic methods. COVID-19 is caused by airborne infection of SARS-CoV-2, and therefore, methods of inactivating the virus efficiently and safely are crucial for reducing the risk of airborne infection. In this regard, the effect of plasma discharge on the infectivity of the coronaviruses mouse hepatitis virus (MHV) and SARS-CoV-2 was tested. Plasma discharge efficiently reduced the infectivity of both coronaviruses. Treatment of SARS-CoV-2 in culture medium with a plasma discharge resulted in 95.17% viral inactivation after plasma irradiation after 1 hour (hr), 99.54% inactivation after 2 hrs and 99.93% inactivation after 3 hrs. Similar results were obtained for MHV. The results indicated that plasma discharge effectively and safely inactivated the airborne coronaviruses and may be useful in minimizing the risk of airborne infection of SARS-CoV-2.

## Introduction

Coronavirus disease 2019 (COVID-19) is characterized by fever, cough, myalgia to severe acute respiratory distress, and multiorgan dysfunction in the case of severe cases, and the clinical manifestations are similar to severe acute respiratory syndrome (SARS) and middle east respiratory syndrome (MERS). COVID-19 was first identified in December 2019 in Wuhan, China, and then spread throughout the world [1–3]. The World Health Organization declared the COVID-19 outbreak to be a pandemic in March 2020, and since then the numbers of infections around the world rose, reaching more than 25 million infections and over 850,000 deaths by 15 September 2020. COVID-19 is caused by a novel coronavirus, SARS coronavirus-2 (SARS-CoV-2), belonging to the genus *Betacoronavirus, family Coronaviridae* [1–4]. Since coronaviruses are enveloped RNA viruses, many disinfectants such as a variety of detergents and alcohols, such as ethanol and 2-propanol, have been shown to inactivate SARS-CoV-2 infectivity [5–23].

SARS-CoV-2 transmits from human to human mainly via airborne routes such as droplets and aerosols [24]; however, the virus also infects by direct contact with virus-contaminated surfaces [25]. In fact, the virus has been shown to be relatively stable on plastic and stainless steel surfaces and on surgical masks for up to 7 days [26–28]. To reduce or minimize the risk of airborne infection of SARS-CoV-2, it is important to disinfect the virus in aerosols and/or droplet nuclei.

Since non-thermal plasmas generated at atmospheric pressure produce active molecules that cause various chemical reactions at room temperature, they are being investigated for use as a process technology with high energy efficiency [29]. Among such plasmas, a “streamer” is a discharge mode in which a wide plasma region can be formed between the electrodes [30], and thus it is the most efficient and capable of generating a large number of active molecules. If viruses are exposed to the active molecules generated from streamers, it is expected that proteins on the surfaces of the viruses will be denatured by chemical reactions and their viral activity will be lost. In this report, we analyzed this inactivating effect by exposing coronaviruses to streamers.

## Materials and Methods

### Cells

Mouse astrocytoma-derived DBT cells were obtained from the Japanese Collection of Research Bioresources (https://cellbank.nibiohn.go.jp/english/) (JCRB) and cultured in Eagle’s medium (MEM, Fujifilm Woko Chemicals, Japan) supplemented with 5% heat-inactivated fetal bovine serum (FBS, Thermo Fisher Scientific, Rockford, IL), 1% (w/v) tryptose phosphate broth (Sigma-Aldrich, St. Louis, MO) and antibiotics (10 units/mL penicillin and 10 μg/mL streptomycin, Fujifilm Wako Chemicals).

Vero E6 cells and Vero E6/TMRPSS2 cells obtained from JCRB Cell Bank under no. JCRB1819, Japan were cultured at 37 °C in Dulbecco’s modified Eagle’s medium (DMEM, Nacalai Tesque, Japan) supplemented with 5% heat-inactivated fetal bovine serum (FBS, Thermo Fisher Scientific, Rockford, IL) and antibiotics (10 units/mL penicillin and 10 μg/mL streptomycin, Thermo Fisher Scientific).

### Viruses

MHV, strain A59, was obtained from the American Type Culture Collection (Summit Pharmaceuticals International, Japan) and propagated in DBT cells in DMEM/Ham’s F-12 medium (Fujifilm Wako Chemicals) supplemented with 10% FBS and antibiotics.

SARS-CoV-2, strain AI/I-004/2020, was obtained from the National Institute of Infectious Diseases, Tokyo, Japan and propagated in Vero E6 cells in DMEM supplemented with 1% FBS and antibiotics.

### Plasma irradiation

A plasma discharge apparatus was used to irradiate the viruses in the media. Part of the plasma discharge device was mounted on one of a series of streamer air purifier products of Daikin Industries, Ltd., and a high voltage was applied to a discharge device having multiple pairs of discharge electrodes to cause a streamer discharge. In the experiment, the virus was placed in a 6-well plate (0.5 mL/well) without a top cover (Product No. 3335, Corning, NY), and the discharge device was placed 10 cm above the plate. The entire experimental system was placed in an acrylic container having an internal volume of 30.9 L (24 cm × 46 cm × 28 cm) and sealed. After applying a voltage of 6-7 kV between the electrodes to cause a current of 55 μA and exposing the virus to the plasma at 25 °C for 0, 1, 2, and 3 hours, respectively, then the virus was collected and subjected to virus infectivity titration.

### Virus titration

Titration of MHV was performed as described previously. Briefly, DBT cells seeded on 60 mm dishes were inoculated with serially 10-fold diluted samples and incubated at 37 °C for 1 hr in a 5% CO_2_ incubator. Then the inoculum was replaced with an agarose medium consisting of MEM supplemented with 2% FBS, 1% (w/v) tryptose phosphate broth, 1% agarose and antibiotics and then cultured for 2 days at 37 °C in 5% CO_2_. The plaques caused by MHV infection were stained by an overlay of agarose medium containing 0.01% neutral red (Sigma-Aldrich). The infectious titer of MHV was expressed as a plaque-forming unit (pfu)/mL. The infectious titer of SARS-CoV-2 was obtained as a 50% tissue culture infectious dose (TCID_50_/mL). The Vero E6/TMPRSS2 cells cultured in 96-well microplates were inoculated with serially 10-fold diluted samples (each 6-wells/dilution) and incubated for 4 days at 37 °C in 5% CO_2_. The numbers of wells showing cytopathic effect (CPE) at each dilution were counted, and TCID_50_/mL was calculated by the Reed–Muench method.

### TaqMan RT-PCR

RNA copies upon plasma irradiation of SARS-CoV-2 were quantitatively determined by using the TaqMan RT-PCR method as described previously [31], with slight modification. Briefly, the virus specimens before and after plasma irradiation were treated with Buffer AVL containing carrier RNA, and RNA was extracted according to the manufacturer’s instructions for QIAmp Viral RNA Mini Kit (Qiagen, Hilden, Germany).

TaqMan RT-PCR was performed with the N2 primer/probe set and BigDye Terminator v3.1 Cycle Sequencing Kit (Thermo Fischer Scientific) [31] using the QuantStudio 5 system (Thermo Fisher Scientific, Rockford, IL).

### Biosafety

Experiments using infectious MHV and SARS-CoV-2 were conducted in a biosafety level (BSL)-2 laboratory in The University of Tokyo and a BSL-3 laboratory in Okayama University of Science, respectively.

## Results

### Inactivation of MHV upon plasma irradiation

We examined the inactivation of MHV after irradiation with the plasma streamer. The infectivity of the virus in the medium after 1, 2, and 3 hrs plasma irradiation was measured and expressed as pfu/mL (Table 1). In mock irradiation (control), the virus infectivity was determined to be 5.26 × 10^7^ pfu/mL, 5.64 × 10^7^ pfu/mL, 5.45 × 10^7^ pfu/mL and 3.56 × 10^7^ pfu/mL after 0, 1, 2 and 3 hrs, respectively. The virus infectivity upon plasma stream irradiation was 2.06 × 10^7^ pfu/mL, 1.57 × 10^6^ pfu/mL, and 5.42 × 10^4^ pfu/mL after irradiation for 1, 2, and 3 hrs, respectively. When compared to the titers of corresponding times of the mock irradiation, the reductions in the virus titers upon plasma irradiation were 63.44%, 97.12%, and 99.85% after 1, 2, and 3 hrs irradiation, respectively.

**Table 1.** Effect of plasma irradiation on the infectivity of MHV.

|  | PFU/mL |  |  |  |  | % reduction to control |  |  |  |  |
| --- | --- | --- | --- | --- | --- | --- | --- | --- | --- | --- |
| experiment | 1 | 2 | 3 | 4 | average | 1 | 2 | 3 | 4 | average |
| control 0hr | $5.62 \times 10^7$ | $5.62 \times 10^7$ | $4.90 \times 10^7$ | $4.90 \times 10^7$ | $5.26 \times 10^7$ | | | | | |
| control 1hr | $6.31 \times 10^7$ | $8.32 \times 10^7$ | $3.16 \times 10^7$ | $4.79 \times 10^7$ | $5.64 \times 10^7$ | | | | | |
| control 2hr | $3.47 \times 10^7$ | $9.12 \times 10^7$ | $1.07 \times 10^7$ | $8.13 \times 10^7$ | $5.45 \times 10^7$ | | | | | |
| control 3hr | $5.75 \times 10^7$ | $1.58 \times 10^7$ | $3.02 \times 10^6$ | $6.61 \times 10^7$ | $3.56 \times 10^7$ | | | | | |
| plasma 1hr | $2.00 \times 10^7$ | $5.62 \times 10^7$ | $3.39 \times 10^6$ | $2.95 \times 10^6$ | $2.06 \times 10^7$ | 68.38% | 32.39% | 89.29% | 93.83% | 63.44% |
| plasma 2hr | $9.55 \times 10^5$ | $4.90 \times 10^6$ | $9.33 \times 10^4$ | $3.24 \times 10^5$ | $1.57 \times 10^6$ | 97.25% | 94.63% | 99.13% | 99.60% | 97.12% |
| plasma 3hr | $6.61 \times 10^4$ | $1.15 \times 10^5$ | $4.17 \times 10^3$ | $3.16 \times 10^4$ | $5.42 \times 10^4$ | 99.89% | 99.28% | 99.86% | 99.95% | 99.85% |

### Inactivation of SARS-CoV-2 upon plasma irradiation

We examined the inactivation of SASR-CoV-2 after irradiation with the plasma streamer. The infectivity of the virus in the medium after 1, 2, and 3 hrs plasma irradiation was measured and expressed as TCID_50_/mL (Table 2). In mock irradiation (control), the virus infectivity was 2.23 × 10^7^ TCID_50_/mL, 2.17 × 10^7^ TCID_50_/mL, 1.10 × 10^7^ TCID_50_/mL, and 9.96 × 10^6^ TCID_50_/mL after 0, 1, 2, and 3 hrs, respectively. The virus infectivity upon plasma stream irradiation was 1.05 × 10^6^ TCID_50_/mL, 5.01 × 10^4^ TCID_50_/mL, and 6.53 × 10^3^ TCID_50_/mL after irradiation for 1, 2, and 3 hrs, respectively. When compared to the titers of the corresponding times of the mock irradiation, the reductions of the virus titers upon plasma irradiation were 95.17%, 99.54%, and 99.93% after 1, 2, and 3 hrs irradiation, respectively (Table 2).

**Table 2.** Effect of plasma irradiation on the infectivity of SARS-CoV-2

|  | TCID<sub>50</sub>/mL |  |  |  |  | %reduction to control |  |  |  |  |
| --- | --- | --- | --- | --- | --- | --- | --- | --- | --- | --- |
| experiment | 1 | 2 | 3 | 4 | average | 1 | 2 | 3 | 4 | average |
| control 0hr | $2.50 \times 10^7$ | $1.41 \times 10^7$ | $2.50 \times 10^7$ | $2.50 \times 10^7$ | $2.23 \times 10^7$ | | | | | |
| control 1hr | $1.41 \times 10^7$ | $1.41 \times 10^7$ | $1.41 \times 10^7$ | $4.45 \times 10^7$ | $2.17 \times 10^7$ | | | | | |
| control 2hr | $1.41 \times 10^7$ | $1.41 \times 10^7$ | $7.94 \times 10^6$ | $7.94 \times 10^6$ | $1.10 \times 10^7$ | | | | | |
| control 3hr | $1.00 \times 10^7$ | $1.41 \times 10^7$ | $7.94 \times 10^6$ | $7.94 \times 10^6$ | $9.96 \times 10^6$ | | | | | |
| plasma 1hr | $7.91 \times 10^5$ | $9.95 \times 10^5$ | $9.95 \times 10^5$ | $1.41 \times 10^6$ | $1.05 \times 10^6$ | 94.38% | 92.92% | 92.92% | 96.84% | 95.17% |
| plasma 2hr | $4.45 \times 10^4$ | $1.41 \times 10^4$ | $7.91 \times 10^4$ | $6.28 \times 10^4$ | $5.01 \times 10^4$ | 99.68% | 99.90% | 99.00% | 99.21% | 99.54% |
| plasma 3hr | $2.50 \times 10^3$ | $5.78 \times 10^3$ | $9.95 \times 10^3$ | $7.91 \times 10^3$ | $6.53 \times 10^3$ | 99.97% | 99.96% | 99.87% | 99.90% | 99.93% |

### Effect of plasma irradiation on SARS-CoV-2 RNA copies of SARS-CoV-2 in the medium

We examined the effect of the number of RNA copies of SASR-CoV-2 after irradiation with the plasma streamer. The RNA of the virus in the medium after 1, 2, and 3 hrs of mock irradiation or plasma streamer irradiation in experiment 1 described above was extracted. The RNA was extracted from the plasma-irradiated and mock-irradiated viruses, and then the RNA was diluted 1,000-fold and subjected to TaqMan RT-PCR. In the mock irradiation, the numbers of virus RNA copies were calculated to be slightly reduced from those of the 0 hr treated virus of 1.1 × 10^11^ RNA copies/mL to 1.0 × 10^11^ RNA copies/mL, 9.1 × 10^10^ RNA copies/mL, and 8.3 × 10^10^ RNA copies/mL after mock irradiation for 1, 2 and 3 hrs, respectively. These reductions in the RNA copies/mL compared to the 0 hr treated virus were 8.3%, 17.2%, and 21.4% after 1, 2, and 3 hrs, respectively. The numbers of virus RNA copies upon plasma stream irradiation were 1.7 × 10^10^ RNA copies/mL, 2.9 × 10^9^ RNA copies/mL, and 6.9 × 10^8^ RNA copies/mL after irradiation for 1, 2, and 3 hrs, respectively. When compared to the RNA copies of the corresponding times with the mock irradiation, the reductions of the virus RNA copies upon plasma irradiation were 82.94%, 96.79%, and 99.17% after 1, 2, and 3 hrs irradiation, respectively (Table 3).

**Table 3.** Effect of plasma irradiation on the RNA copies of SASR-CoV-2

|  | experiment | RNA copies / mL<br>virus | average<br>RNA copies /mL | % reduction to<br>the control |
| --- | --- | --- | --- | --- |
| control 0hr | #1 | $1.08 \times 10^{11}$ | $1.10 \times 10^{11}$ | |
| | #2 | $1.11 \times 10^{11}$ | | |
| control 1hr | #1 | $9.96 \times 10^{10}$ | $1.00 \times 10^{11}$ | |
| | #2 | $1.01 \times 10^{11}$ | | |
| control 2hr | #1 | $8.72 \times 10^{10}$ | $9.07 \times 10^{10}$ | |
| | #2 | $9.43 \times 10^{10}$ | | |
| control 3hr | #1 | $8.37 \times 10^{10}$ | $8.31 \times 10^{10}$ | |
| | #2 | $8.26 \times 10^{10}$ | | |
| plasma 1hr | #1 | $1.71 \times 10^{10}$ | $1.71 \times 10^{10}$ | 82.94% |
| | #2 | $1.71 \times 10^{10}$ | | |
| plasma 2hr | #1 | $2.96 \times 10^9$ | $2.91 \times 10^9$ | 96.79% |
| | #2 | $2.86 \times 10^9$ | | |
| plasma 3hr | #1 | $6.73 \times 10^8$ | $6.90 \times 10^8$ | 99.17% |
| | #2 | $7.08 \times 10^8$ | | |

## Discussion

The stability of coronaviruses, including SARS-CoV-2, in different environmental conditions has been reported. A variety of commercially available disinfectants have been shown to be effective in inactivating SARS-CoV-2. However, the virus has been shown to be relatively stable in dried form, especially on stainless plastic surfaces and on the outer layers of masks, and infectious virus has been recovered after up to 4 to 7 days. The virus has been shown to be more stable in liquid form. Coronaviruses like MHV and SARS-CoV-2 are thought to infect susceptible animals and humans by aerosols and direct contact. To reduce or minimize the risk of airborne infection of SARS-CoV-2, it is important to disinfect the virus in aerosols and/or droplet nuclei. Recently, studies on the inactivation of viruses, bacteria, and fungi by plasma irradiation have been reported. However, the mechanism of inactivation of microorganisms by plasma irradiation is not fully understood. Streamer discharge technology, which is a type of plasma discharge, produces various active substances by the collision of electrons moving at high speed in the plasma region with molecules such as nitrogen, oxygen, and water vapor in the atmosphere. For example, metastable nitrogen is considered to play an important role in various chained chemical reactions due to its high energy level and long life [32, 33].

Other studies have examined methods of exposing microorganisms directly to plasma [34]. According to these methods, high-energy electrons and short-lived active species, which are always present in large quantities in the plasma space, can be used. Therefore, the inactivation efficiency is increased, and the time required is shortened. However, this method is suitable for inactivating small areas only and is not suitable for processing large areas. Thus, it is not considered to inactivate microorganisms efficiently when their concentrations are low.

In the present study, the inactivation rates of two coronaviruses, MHV and SARS-CoV-2, were virtually identical, indicating that the mechanisms of inactivation were the same. The decay rate of SARS-CoV-2 RNAs determined by RT-qPCR was slightly slower compared to that determined by SARS-CoV-2 infectivity. The RT-qPCR used in the present study detected 138 bases of the virus RNA; thus, the decay of the RNA measured by the RT-qPCR detected disruption of the virus RNA into fewer than 138 bases. However, virus infectivity will be lost even if there is only partial damage to the virus RNA with a size of 29,903 bases caused by plasma irradiation because an intact virus genome is necessary for the virus to replicate in the infected cells. In this regard, it is likely that SARS-CoV-2 inactivation based on RT-qPCR was underestimated compared to that based on the infectivity assay. Thus, the virus infectivity was reduced or lost by both a loss of virus protein functions and a disruption of viral RNAs.

In the present study, various active molecules are generated in the plasma region of a streamer in space and exposed to a target away from the plasma. Although the time required for inactivation is slightly longer [34], this form of application is suitable for inactivating large spaces and large amounts of air. In this regard, a quantitative measurement of such active molecules is a difficult technique that requires sophisticated equipment and skills. Further progress is necessary to clarify the mechanism of virus inactivation in the future.

In conclusion, a plasma streamer was shown to be effective in inactivating the coronaviruses MHV and SARS-CoV-2.

